# Distinct Neutralizing Antibody Escape of SARS-CoV-2 Omicron Subvariants BQ.1, BQ.1.1, BA.4.6, BF.7 and BA.2.75.2

**DOI:** 10.1101/2022.10.19.512891

**Authors:** Panke Qu, John P. Evans, Julia Faraone, Yi-Min Zheng, Claire Carlin, Mirela Anghelina, Patrick Stevens, Soledad Fernandez, Daniel Jones, Gerard Lozanski, Ashish Panchal, Linda J. Saif, Eugene M. Oltz, Kai Xu, Richard J. Gumina, Shan-Lu Liu

## Abstract

Continued evolution of SARS-CoV-2 has led to the emergence of several new Omicron subvariants, including BQ.1, BQ. 1.1, BA.4.6, BF.7 and BA.2.75.2. Here we examine the neutralization resistance of these subvariants, as well as their ancestral BA.4/5, BA.2.75 and D614G variants, against sera from 3-dose vaccinated health care workers, hospitalized BA.1-wave patients, and BA.5-wave patients. We found enhanced neutralization resistance in all new subvariants, especially the BQ.1 and BQ.1.1 subvariants driven by a key N460K mutation, and to a lesser extent, R346T and K444T mutations, as well as the BA.2.75.2 subvariant driven largely by its F486S mutation. The BQ.1 and BQ.1.1 subvariants also exhibited enhanced fusogenicity and S processing dictated by the N460K mutation. Interestingly, the BA.2.75.2 subvariant saw an enhancement by the F486S mutation and a reduction by the D1199N mutation to its fusogenicity and S processing, resulting in minimal overall change. Molecular modelling revealed the mechanisms of receptor-binding and non-receptor binding monoclonal antibody-mediated immune evasion by R346T, K444T, F486S and D1199N mutations. Altogether, these findings shed light on the concerning evolution of newly emerging SARS-CoV-2 Omicron subvariants.

## Introduction

Since its emergence in late 2021, the Omicron variant of severe acute respiratory syndrome coronavirus 2 (SARS-CoV-2) has produced numerous subvariants that continue to erode vaccine- and infection-induced immunity, and alter virus biology (Evans et al., 2022; Kurhade et al., 2022; Qu et al., 2022a; Qu et al., 2022b; Xia et al., 2022b; Yamasoba et al., 2022; Yu et al., 2022). The initial BA.1 Omicron subvariant spurned a large wave of coronavirus disease 2019 (COVID-19) cases and exhibited strong immune escape from 2-mRNA vaccine dose-induced immunity that was recovered by booster mRNA vaccine administration (Abu-Raddad et al., 2022; Cao et al., 2022b; Cerutti et al., 2022; Evans et al., 2022; Gruell et al., 2022; Liu et al., 2022; Xia et al., 2022a; Zou et al., 2022). In addition, the BA.1 subvariant exhibited reduced cell-cell fusogenicity, impaired replication in lower airway epithelial cells, as well as altered entry route preference (Barut et al., 2022; Cui et al., 2022; Meng et al., 2022; Shuai et al., 2022; Suzuki et al., 2022; Wang et al., 2022a). These features correlated with a reduced replication capacity of Omicron in lung tissues, enhanced nasopharyngeal tropism, and decreased pathogenicity *in vivo* (Barut et al., 2022; McMahan et al., 2022; Shuai et al., 2022; Su et al., 2022b; Suzuki et al., 2022). Importantly, these characteristics were largely maintained by subsequent Omicron subvariants.

The BA.2 subvariant overtook BA.1 due to its slightly enhanced transmissibility and immune evasion, with an ability to reinfect individuals who were previously infected with BA.1 (Iketani et al., 2022; Lyngse et al., 2022; Stegger et al., 2022). From BA.2, several subvariants emerged in quick succession, often with concurrent circulations; these included the BA.4 and BA.5 subvariants (bearing identical S proteins, referred to as BA.4/5 hereafter) that next rose to dominance and exhibited further immune escape (Cao et al., 2022c; Hachmann et al., 2022; Khan et al., 2022; Kimura et al., 2022; Qu et al., 2022b; Tuekprakhon et al., 2022; Wang et al., 2022b). In addition, BA.2 gave rise to the BA.2.75 subvariant, which is currently increasing in proportion of COVID-19 cases (Centers for Disease Control and Prevention, 2022), but does not exhibit as substantial immune escape compared to BA.4/5 (Cao et al., 2022a; Qu et al., 2022a; Saito et al.; Wang et al., 2022c). The BA.4/5 and BA.2.75 subvariants have driven further diversification of the circulating SARS-CoV-2, with the emergence of several additional subvariants including the BA.4.6, BF.7, BQ.1, and BQ.1.1 (derived from BA.4/5), as well as BA.2.75.2 (derived from BA.2.75). These new subvariants are currently increasing in frequency (Centers for Disease Control and Prevention, 2022; Iacobucci, 2022) and may be the next major dominant Omicron subvariant.

The extent of immune evasion and functional alterations to the spike protein (S) in these emerging Omicron subvariants remains unclear. To address this, we examine the resistance of the BA.4.6, BF.7, BQ.1, BQ.1.1, and BA.2.75.2 subvariants to neutralization by serum from recipients of 3 mRNA vaccine doses, as well as COVID-19 patients infected with the BA.1 or BA.4/5 variants. We observe strong neutralization resistance in the BQ.1 and BQ.1.1 subvariants driven largely by their key N460K mutation, as well as in the BA.2.75.2 subvariant driven largely by its singature F486S mutation. We further examine the fusogenicity and processing of the subvariant S proteins and observe enhanced fusogenicity and S processing in the BA.4/5-derived subvariants driven largely by the N460K mutation, but comparable fusogenicity and S processing in BA.2.75-derived BA.2.75.2 subvariant modulated by its F485S and D1199N mutations. Finally, structural modeling showed that the F486S mutation reduced binding affinity for both the ACE2 receptor and class I and II antibodies, while the R346T and K444T mutations are likely responsible for evasion of class III antibody recognition.

## Results

### BQ.1, BQ1.1, and BA.2.75.2 exhibit potent neutralization resistance

To examine the neutralization resistance of emerging Omicron subvariants, we utilized our previously reported pseudotyped lentivirus neutralization assay (Zeng et al., 2020). Lentivirus pseudotyped with S from each of the critical Omicron subvariants (**Fig. 1A**), as well as from the prototype D614G variant were produced. All Omicron subvariant-pseudotyped viruses exhibited modestly enhanced infectivity in HEK293T-ACE2 cells over the D614G variant, with the exception of BA.2.75.2 (**Fig. 1B**). Additionally, all Omicron subvariants exhibited comparably poor infectivity in lung-derived CaLu-3 cells (**Fig. 1C**) compared to D614G, consistent with prior Omicron subvariants, and the weak lung tropism observed for the Omicron variant (Barut et al., 2022; Meng et al., 2022; Shuai et al., 2022; Wang et al., 2022a).

**Figure 1:**
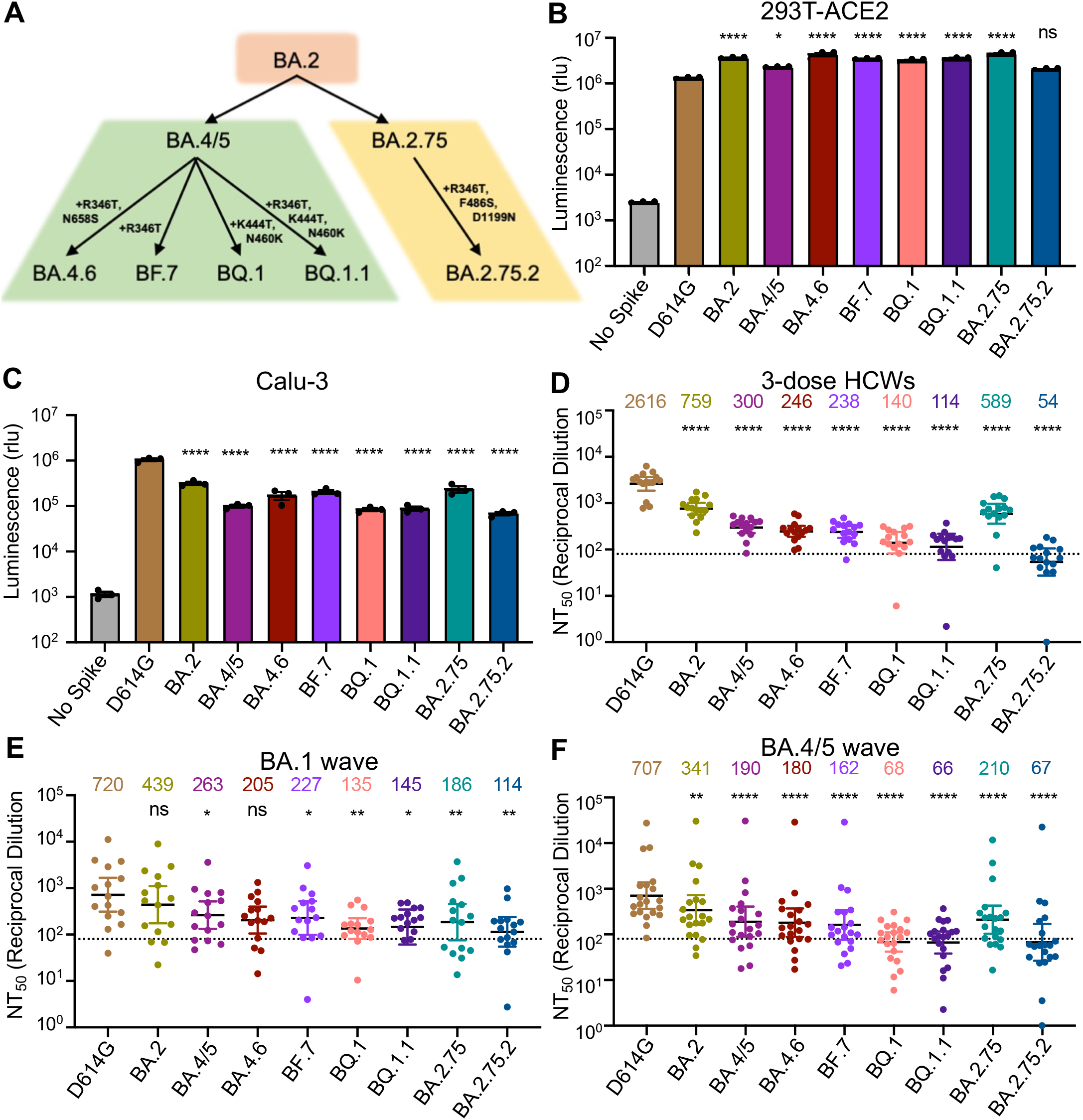
Omicron subvariants, especially BQ.1, BQ.1.1, and BA.2.75.2, exhibit strong neutralization resistance. (**A**) Displayed is a schematic of SARS-CoV-2 Omicron subvariant evolution indicating the mutations acquired by the BA.4.6, BF.7, BQ.1, BQ.1.1, and BA.2.75.2 subvariants. (**B-C**) Infectivity of lentivirus pseudotyped with the indicated S constructs in HEK293T-ACE2 cells, n = 3 (B), or lung-derived CaLu-3 cells, n = 3 (C). Bars represent means +/− standard error. Significance relative to D614G was determined by one-way ANOVA with Bonferroni’s multiple testing correction. P values are represented as ns for p ≥ 0.05, *p < 0.05, and ****p < 0.0001, respectively. (**D-F**) Neutralizing antibody tiers against lentivirus pseudotyped with S from the indicated SARS-CoV-2 variants were determined for sera from health care workers (HCWs) (n = 15) who received a single homologous monovalent Moderna mRNA-1273 (n =3) or Pfizer/BioNTech BNT162b2 (n =12) mRNA booster vaccination (D), for sera from BA.1-wave hospitalized COVID-19 patients (n = 15) (E), and for sera from BA.4/5-wave SARS-CoV-2 infected Columbus, Ohio first responders and household contacts (n = 20) (F). Bars represent geometric means with 95% confidence intervals. Significance relative to D614G was determined by one-way repeated measures ANOVA with Bonferroni’s multiple testing correction. P values are displayed as ns for p ≥ 0.05, *p < 0.05, **p < 0.01, and ****p < 0.0001.

We next examined the resistance of these emerging Omicron subvariants to sera from health care workers (HCWs) collected 2-13 weeks after vaccination with a homologous booster dose of monovalent Moderna mRNA-1273 (n =3) or Pfizer/BioNTech BNT162b2 vaccine (n =12). We observed potent neutralization resistance among all Omicron subvariants compared to ancestral D614G (**Fig. 1D** and **Fig. S1A**). Specifically, compared to D614G, the BA.4.6, BF.7, BQ.1, and BQ.1.1 subvariants exhibited a 10.6-fold (p < 0.0001), 11.0-fold (p < 0.0001), 18.7-fold (p < 0.0001), and 22.9-fold (p < 0.0001) lower neutralization sensitivity, respectively, while BA.4/5 exhibited an 8.7-fold (p < 0.0001) lower neutralization sensitivity than D614G (**Fig. 1D**). Similarly, compared to D614G, the BA.2.75.2 subvariant exhibited a 48.4-fold (p < 0.0001) lower neutralization sensitivity while BA.2.75 exhibited a 4.4-fold (p < 0.0001) lower neutralization sensitivity than D614G (**Fig. 1D**). These data indicate further neutralization escape in emerging Omicron subvariants, with BA.Q.1, BA.Q.1.1, and BA.2.75.2, especially the latter, showing the most substantial neutralization resistance.

We also examined the resistance of Omicron subvariants to neutralization by sera from hospitalized COVID-19 patients (n =15) infected during the BA.1 wave of the pandemic. Despite exposure to an Omicron antigen, these patients displayed a remarkably similar neutralization resistance pattern to the boosted HCWs. Specifically, compared to D614G, the BA.4.6, BF.7, BQ.1, and BQ.1.1 subvariants exhibited a 3.5-fold (p>0.05), 3.2-fold (p <0.05), 5.3-fold (p <0.01), and 5.0-fold (p <0.05) lower neutralization sensitivity, respectively, while BA.4/5 exhibited an 2.7-fold (p <0.05) lower neutralization sensitivity than D614G (**Fig. 1E**). Similarly, compared to D614G, the BA.2.75.2 subvariant exhibited a 6.3-fold (p <0.01) lower neutralization sensitivity, while BA.2.75 exhibited a 3.9-fold (p <0.01) lower neutralization resistance (**Fig. 1E**). As would be expected, BA.1 patient sera neutralized BA.2 with a higher efficiency compared to these BA.2-derived subvariants (**Fig. 1E** and **Fig. S1B**).

To determine the breadth of immunity from individuals infected with more recent Omicron subvariants, we next examined sera from Columbus, Ohio first responders and household contacts testing positive for COVID-19 during the BA.4/5 wave of the pandemic, with 11/20 subjects having the infecting variant confirmed as BA.4/5 or a derivative by sequencing. Notably, this cohort exhibited the weakest neutralizing antibody titers of the three groups examined, consistent with milder COVID-19 cases correlating with weaker neutralizing antibody responses (Zeng et al., 2020). Critically, this cohort again displayed a similar pattern of neutralization resistance. Specifically, compared to D614G, the BA.4.6, BF.7, BQ.1, and BQ.1.1 subvariants exhibited a 3.9-fold (p < 0.0001), 4.4-fold (p < 0.0001), 10.4-fold (p < 0.0001), and 10.7-fold (p < 0.0001) lower neutralization sensitivity, respectively, while BA.4/5 exhibited a 3.7-fold (p < 0.0001) lower neutralization sensitivity than D614G (**Fig. 1F**). Additionally, compared to D614G, the BA.2.75.2 subvariant exhibited a 10.6-fold (p < 0.0001) lower neutralization sensitivity whereas BA.2.75 showed a 3.4-fold (p < 0.0001) decreased neutralization sensitivity (**Fig. 1F**). Interestingly, BA.2 exhibited less resistance to sera of BA.4/5-wave infection than BA.4/5, with 2.1-fold reduced neutralization sensitivity compared to D614G (p < 0.01) (**Fig. 1F** and **Fig. S1C**). Together, these results showed that BQ.1, BQ.1.1, and BA.2.75.2 are strongly resistant to neutralization by sera from subjects infected with the recently dominant BA.4/5 variant and suggested that BA.4/5 infection does not offer a broader protection against newly emerging subvariants.

### R346T, K444T, N460K, and F486S represent key neutralization escape mutations

To determine the features governing neutralization resistance, we produced single and double mutant S constructs to probe the contributions of individual amino acid substitutions alone or in combination to neutralization resistance. The panel of mutants exhibited largely similar infectivity compared to their parental BA.4/5 or BA.2.75 variants in HEK293T-ACE2 cells (**Fig. 2A-B**) and Calu-3 cells (**Fig. S2A-B**), although F486S-containing, BA.2.75-derived mutants appear to have modestly reduced titers in Calu-3 cells (**Fig. 2A-B**, and **Fig. S2A-B**). For HCWs who received 3 mRNA vaccine doses introduction of the N460K mutation reduced neutralization sensitivity by an additional 2.6-fold (p < 0.0001) compared to the parental BA.4/5 subvariant, with a similar 2.8-fold (p < 0.0001) reduction for the R346T/N460K double mutant (**Fig. 2C**). Consistent with this finding, the N460K-bearing subvariants BQ.1 and BQ.1.1 showed the strongest neutralization resistance, with a 2.1-fold (p < 0.01) and 2.6-fold (p < 0.01) reduction in neutralization sensitivity compared to BA.4/5 (**Fig. 2C,** and **Fig. S3A**). Though not significant, the other key individual mutations, R346T and K444T, were associated with a milder 20.7% (p =0.0782) and 29.3% (p =0.1886) reduction in neutralization sensitivity, respectively (**Fig. 2C**). Similarly, the N658S mutation did not appear to be strongly associated with neutralization resistance (**Fig. S3A**), with a 23.7 % (p > 0.05) reduction compared to the parental BA.4/5 (**Fig. 2C**). Altogether, these results showed that the N460K mutation, and to a lesser extent R346T, K444T and N658S, are critical for the enhanced resistance to mRNA booster-vaccinated sera in the BQ.1 and BQ.1.1 subvariants.

**Figure 2:**
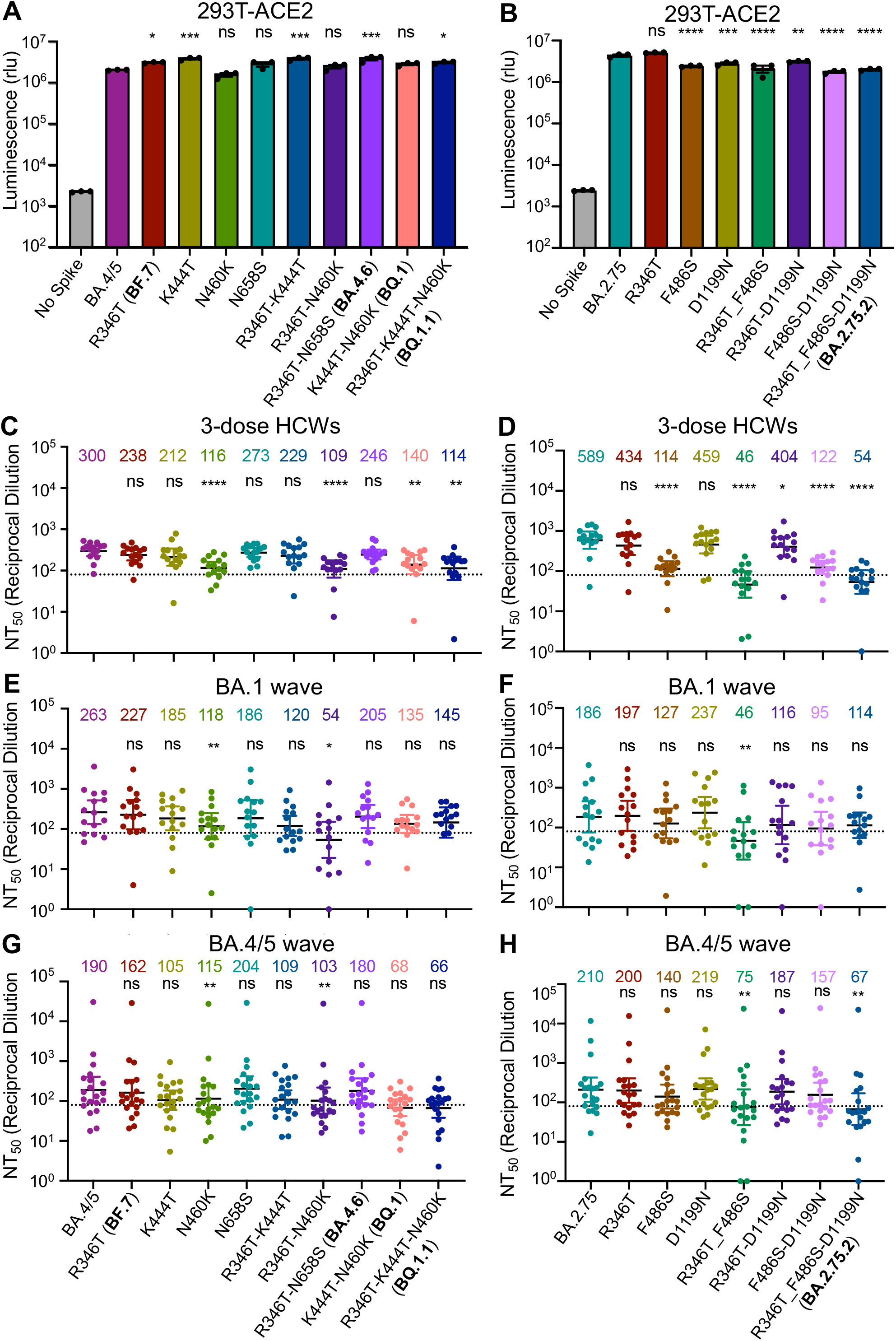
Mutations N460K and F486S, and to a lesser extent, R346T and K444T, drive Omicron subvariant neutralization resistance. (**A-B**) Infectivity of lentivirus pseudotyped with the indicated BA.4/5-derived mutant S constructs, n = 3 (A) or BA.2.75-derived mutant S constructs, n = 3 (B) in HEK293T-ACE2 cells. Bars represent means +/− standard error. Significance relative to BA.4/5 or BA.2.75 was determined by one-way ANOVA with Bonferroni’s multiple testing correction. P values are represented as ns for p ≥ 0.05, *p < 0.05, **p < 0.01, ***p < 0.001, and ****p < 0.0001, respectively. (**C-H**) Neutralizing antibody tiers were determined for sera from health care workers (HCWs) (n = 15) who received a single homologous monovalent Moderna mRNA-1273 (n = 3) or Pfizer/BioNTech BNT162b2 (n = 12) mRNA booster vaccination against lentivirus pseudotyped with S from the BA.4/5-derived mutants (C) and BA.2.75-derived mutants (D); for sera from BA.1-wave hospitalized COVID-19 patients (n = 15) against lentivirus pseudotyped with S from the BA.4/5 derived mutants (E) and BA.2.75 derived mutants (F); and for sera from BA.4/5-wave SARS-CoV-2 infected Columbus, Ohio first responders and household contacts (n = 20) against lentivirus pseudotyped with S from the BA.4/5-derived mutants (G) and BA.2.75-derived mutants (H). Bars represent geometric means with 95% confidence intervals. Significance relative to D614G was determined by one-way repeated measures ANOVA with Bonferroni’s multiple testing correction. P values are displayed as ns for p ≥ 0.05, *p < 0.05, **p < 0.01, and ****p < 0.0001.

For BA.2.75-derived mutants, the introduction of the F486S, R346T/F486S, or F486S/D1199N mutations reduced neutralization sensitivity by 5.2-fold (p < 0.0001), 12.8-fold (p < 0.0001), 4.8-fold (p < 0.0001), respectively, compared to the 10.9-fold (p < 0.0001) reduction seen for the parental BA.2.75.2 subvariant (**Fig. 2D**). However, the introduction of the R346T, D1199N, or R346T/D1199N resulted in only a 26.3% (p = 0.1074), 22.1% (p > 0.05), and 31.4% (p < 0.05) reduction in neutralization sensitivity, respectively (**Fig. 2** and **Fig. S3B**). These results indicated that the F486S mutation is the major driver of enhanced resistance to mRNA booster vaccinee sera in BA.2.75.2.

The pattern of neutralization resistance was similar for the BA. 1-wave hospitalized COVID-19 patient sera. Introduction of the N460K and R346T/N460K mutations into BA.4.5 strongly reduced neutralization sensitivity by 2.2-fold (p < 0.01) and 4.9-fold (p < 0.05), respectively (**Fig. 2E** and **Fig.S3C**). Further, introduction of the R346T/F486S mutations into BA.2.75 reduced neutralization sensitivity by 4.0-fold (p < 0.01) (**Fig. 2F** and **Fig.S3D**). Surprisingly, the F486S mutation alone only resulted in a 31.7% (p = 0.3698) reduction in neutralization sensitivity (**Fig. 2F**).

Consistent with the patterns of HCWs and BA.1-wave patients, neutralization resistance from the BA.4/5-infected COVID-19 patient sera was again largely driven by N460K, F486S, and R346T. Introduction of the N460K and R346T/N460K mutations into BA.4.5 reduced neutralization sensitivity by 1.7-fold (p < 0.01) and 1.8-fold (p < 0.01), respectively (**Fig. 2G** and **Fig. S3E**). Similar to the BA.1-wave patients, introduction of the R346T/F486S mutations into BA.2.75 reduced neutralization sensitivity by 2.8-fold (p < 0.01) (**Fig. 2H** and **Fig.S3F**). Thus, the N460K, F486S, and to a lesser extent R346T, mutations drive neutralization resistance to mRNA vaccinated and boosted, BA.1-infected, and BA.4/5-infected patient sera.

### BQ. 1, and BQ. 1.1 exhibit enhanced fusogenicity and S processing

To examine how alterations to the S protein in these new Omicron subvariants impact function, we examined the ability of the S from these subvariants to mediate cell-cell fusion. Consistent with prior Omicron subvariants (Qu et al., 2022a; Wang et al., 2022a; Zeng et al., 2021), all new subvariants exhibited diminished syncytia formation compared to ancestral D614G (**Fig. 3A-B**), despite comparable levels of cell surface expression of each S construct (**Fig. 3C-D**). Notably, the new Omicron subvariants BA.4.6, BQ.1, and BQ.1.1 exhibited enhanced fusogenicity compared to their parental variant BA.4/5, and similarly, BA.2.75.2 showed an increased fusogenicity relative to BA.2.75 (**Fig. 3C-D**).

**Figure 3:**
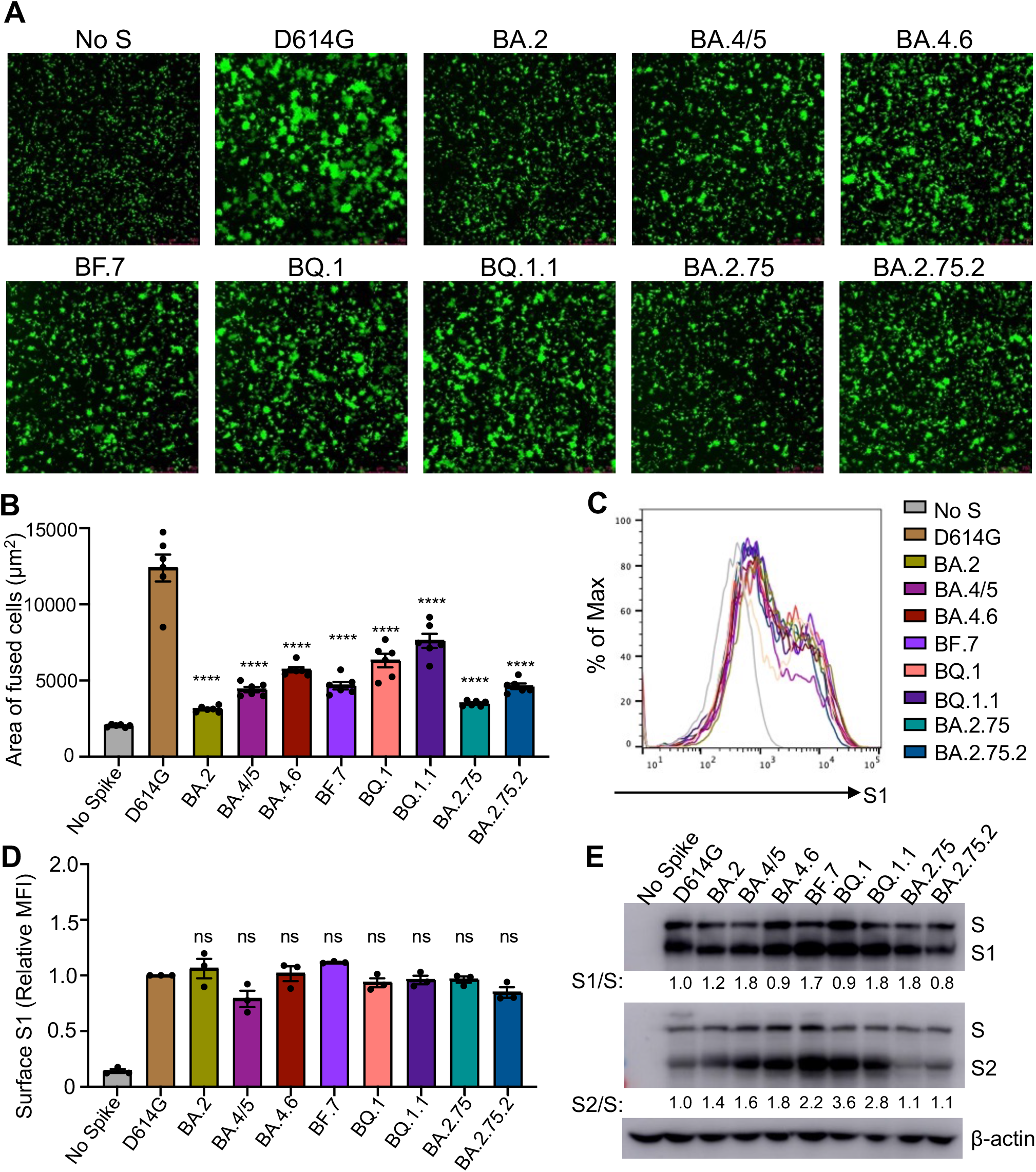
Omicron subvariants, especially BA.4.6, BQ.1 and BQ.1.1, exhibit enhanced fusogenicity and S processing. (**A**) Representative images from syncytia formation assays for S constructs from each of the indicated variants are displayed. The scale bars represent 150 μm. (**B**) Quantification of syncytia formation for the indicated S constructs, n = 6. Bars represent means +/− standard error. Significance relative to D614G was determined by one-way ANOVA with Bonferroni’s multiple testing correction. P values are displayed as ****p < 0.0001. (**C**) Representative histograms of anti-S surface staining in HEK293T cells transfect to express the indicated S constructs. (**D**) Quantification of S surface expression (mean fluorescence intensity) relative to the D614G construct is displayed, n = 3. Bars represent means +/− standard error. Significance relative to D614G was determined by one-way ANOVA with Bonferroni’s multiple testing correction. P values are displayed as ns for p ≥ 0.05. (**E**) S processing is displayed as determined by Western blot probing with anti-S1 and anti-S2 antibodies. The ratio of band intensities for S1 and S as well as S2 and S relative to D614G are displayed.

Given the key role of cellular furin in processing viral S protein into S1 and S2 subunits for SARS-CoV-2 entry (Mykytyn et al., 2021; Peacock et al., 2021), we next examined the proteolytic processing of S in lentivirus producer cells. We saw an enhanced S2/S ratio compared to ancestral D614G in BA.4.6, BF.7, BQ.1, and BQ.1.1 with a 1.8-fold, 2.2-fold, 3.6-fold, and 2.8-fold higher S2/S ratio, respectively, compared to the 1.6-fold higher S2/S ratio in BA.4/5 (**Fig. 3E**). However, compared to BA.4/5, the S1/S ratio of these BA.4/5-derived subvariants was comparable (BF.7 and BQ.1.1) or even reduced (BA.4.6 and BQ.1) (**Fig. 3E**), suggesting possible shedding of S1 into the culture media. BA.2.75 and BA.2.75.2 exhibited minimal difference in S2/S ratio compared to D614G (**Fig. 3E**), but once again, BA.2.75.2 exhibited a decreased S1/S ratio compared to the parental BA.2.75 (**Fig. 3E**), suggesting possibly increased S1 shedding for the BA.2.75.2 subvariant.

### The N460K and F486 mutations enhance fusogenicity while D1199N diminishes fusogenicity

To identify the determinants of enhanced fusogenicity in new Omicron subvariants, we examined S-mediated syncytia formation for single and double S mutants. We observed that introduction of the N460K mutation in BA.4/5 enhanced mean syncytia size by 1.7-fold (p < 0.0001) (**Fig. 4A-B**). Notably all N460K-containing mutants and variants exhibited enhanced syncytia size over BA.4/5 (**Fig. 4A-B**). Further, while the single mutants had no effect, introduction of the R346T/K444T and R346T/N658S mutations slightly enhanced mean syncytia size by 1.4-fold (p < 0.001) and 1.3-fold (p <0.05) over BA.4/5, potentially indicating a synergistic effect likely contributed by R346T (**Fig. 4A-B**). Additionally, while the BA.2.75.2 variant exhibited only modestly enhanced syncytia formation over BA.2.75, i.e., 33.2% (p < 0.0001), the introduction of the F486S or R346T/F486S mutations into BA.2.75 enhanced mean syncytia size by 33.8% (p < 0.0001) and 58.1% (p < 0.0001), respectively, suggesting a critical role of F486S in promoting fusion (Fig. 4C-D). Notably, the introduction of the D1199N mutation into BA.2.75 reduced the mean syncytia size by 17.2% (p < 0.05), revealing opposing effects of the F486S and D1199N mutations on the fusogenicity of BA.2.75.2 (**Fig. 4C-D**). These differences in S-mediated cell-cell fusion occurred despite comparable or even somewhat reduced (especially for BA.2.75.2 mutants) levels of surface expression among these Omicron subvariant S constructs (**Fig. 4E-F, and Fig. S4A-B**).

**Figure 4:**
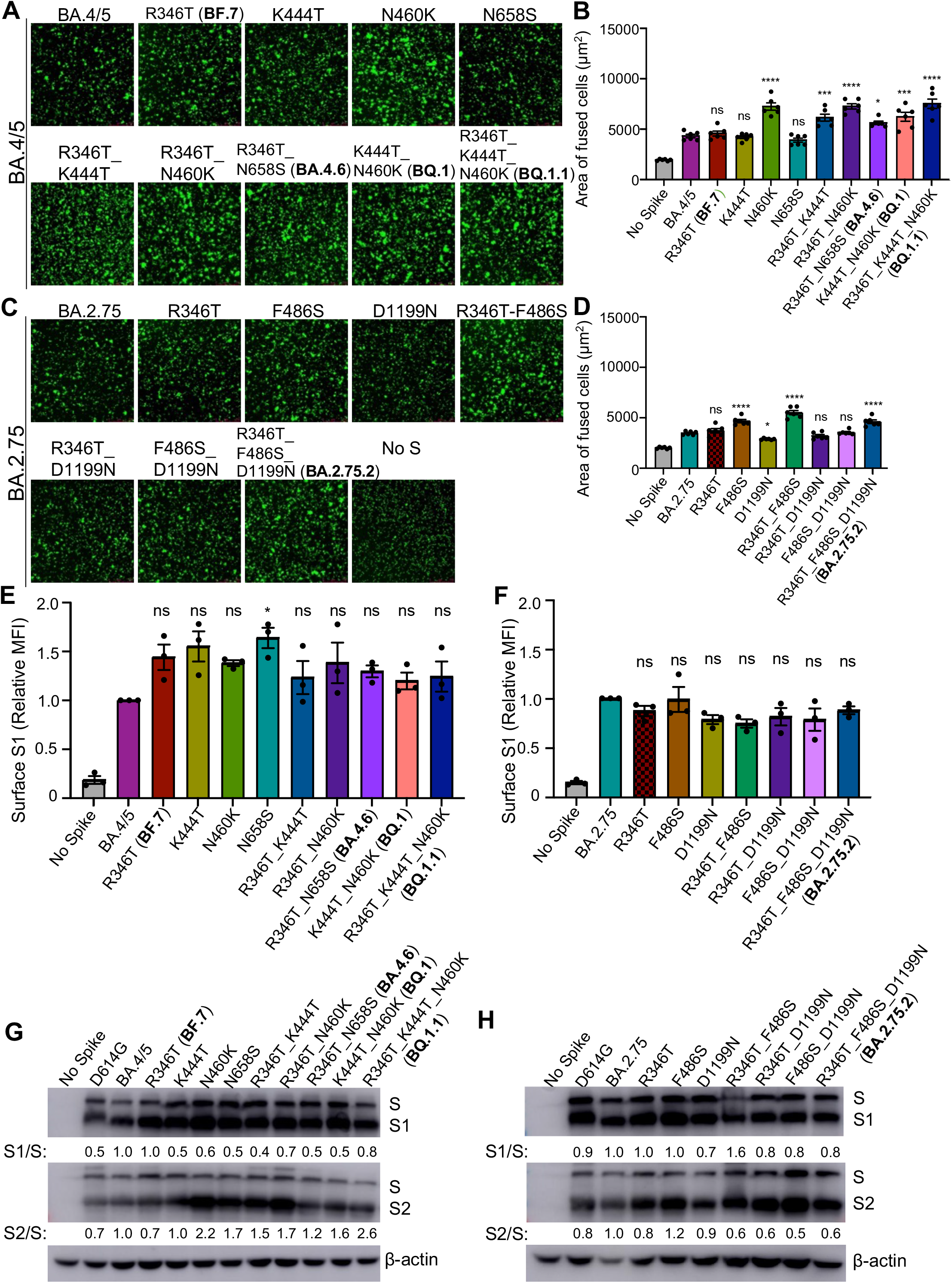
Mutations N460K, N658S, F486S, and D1199N determine the fusogenicity and S processing of Omicron subvariants. (**A**) Representative images from syncytia formation assays for S constructs from each of the indicated BA.4/5-derived mutants are displayed. The scale bars represent 150 μm. (**B**) Quantification of syncytia formation for the indicated BA.4/5 derived S mutants, n = 6. Bars represent means +/− standard error. Significance relative to BA.4/5 was determined by one-way ANOVA with Bonferroni’s multiple testing correction. P values are displayed as ns for p ≥ 0.05, *p < 0.05, ***p < 0.001, and ****p < 0.0001. (**C**) Representative images from syncytia formation assays for S constructs from each of the indicated BA.2.75-derived mutants are displayed. The scale bars represent 150 μm. (**D**) Quantification of syncytia formation for the indicated BA.2.75-derived S mutants, n = 6. Bars represent means +/− standard error. Significance relative to BA.2.75 was determined by one-way ANOVA with Bonferroni’s multiple testing correction. P values are displayed as ns for p ≥ 0.05, *p < 0.05, and ****p < 0.0001. (**E**) Quantification of S surface expression relative to the BA.4/5 construct for the BA.4/5-derived mutants is displayed, n = 3. Bars represent means +/− standard error. Significance relative to BA.4/5 was determined by one-way ANOVA with Bonferroni’s multiple testing correction. P values are displayed as ns for p ≥ 0.05 and *p < 0.05. (**F**) Quantification of S surface expression relative to the BA.2.75 construct for the BA.2.75 derived mutants is displayed, n = 3. Bars represent means +/− standard error. Significance relative to BA.4/5 was determined by one-way ANOVA with Bonferroni’s multiple testing correction. P values are displayed as ns for p ≥ 0.05. (**G**) S processing for the BA.4/5-derived mutants as well as ancestral D614G is displayed, which was determined by Western blot probing with anti-S1, anti-S2, and anti-β-actin antibodies. The ratio of band intensities for S1 and S as well as S2 and S relative to BA.4/5 are displayed. (**H**) S processing for the BA.2.75-derived mutants as well as ancestral D614G is displayed as determined by Western blot probing with anti-S1, anti-S2, and anti-β-actin antibodies. The ratio of band intensities for S1 and S as well as S2 and S relative to BA.2.75 are displayed.

We next sought to determine the impact of the subvariant determining mutations on S processing. For the BA.4/5-derived subvariants, introduction of the N460K, N658S, R346T/K444T, and R346T/N460K mutations exhibited 2.2-fold, 1.7-fold, 1.5-fold, and 1.7-fold higher S2/S ratios, largely consistent with those mutants exhibiting enhanced fusogenicity and S processing, especially for the BQ.1 and BQ.1.1 subvariants (**Fig. 4G**). Remarkably, the S1/S ratio for almost all mutants examined, except the R346T mutation, was decreased compared to the BA.4/5 subvariant (**Fig. 4G**), suggesting an enhanced S1 shedding in these BA.4/5 derived mutants, which is consistent with increased fusion. For BA.2.75-derived mutants, the impacts on S2/S and S1/S ratios were fairly modest, with most exhibiting reductions relative to the parental BA.2.75, consistent with the more modest impact on cell-cell fusion of these variants (**Fig. 4H**). Notably, the introduction of the F486S mutation increased the S2/S ratio by 1.2-fold but had no effect on S1/S ratio compared to BA.2.75 (**Fig. 4H**).

### Structural modeling reveals mechanism of mutation-mediated antibody evasion and alteration in receptor utilization

To further understand the functional impact of the mutations in these Omicron subvariants, we performed homology modeling-based structural analyses. R346 and K444 are located outside of SARS-CoV2 receptor binding motif (RBM) and are within the epitope of class III neutralizing antibodies. Structural analysis indicated an interference of antibody recognition introduced by R346S (**Fig. 5A**) and K444T (**Fig. 5B**), where hydrogen bonds and a salt-bridge can be abolished. In contrast, F486 is located within the RBM and is a key residue for binding to both ACE2 receptor and neutralizing antibodies; F486 interacts hydrophobically with M82 and Y83 on ACE2 (**Fig. 5C**), whereas the F486S mutation negatively impacts the interaction with ACE2, as well as binding by some monoclonal antibodies in class I and III categories. Further structural analysis showed that residue D1199 is located in the heptad repeat 2 (HR2) domain on a solvent-accessible surface close to the transmembrane domain or membrane (**Fig. 5D**). Electrostatic surface potential (**Fig. 5C inset**) of this region reveals a strong negative surface charge, which repulses the negatively charged membrane and could help keep the spike in an up-right position. However, the D1199N mutation in the S2 subunit would reduce the electrostatic repulsion, resulting in a more tilted spike and rendering its less efficient processing and receptor utilization.

**Figure 5:**
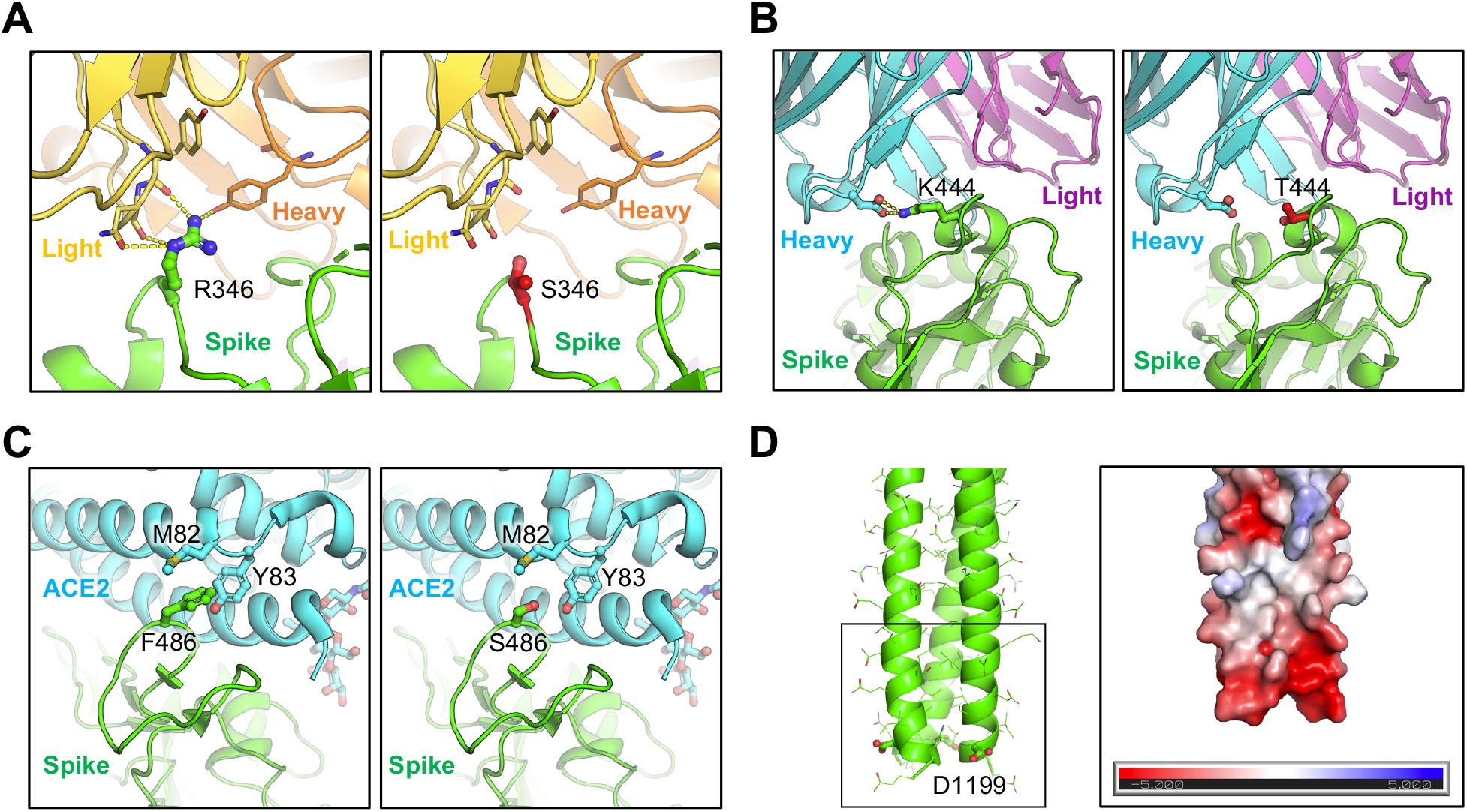
Homology modeling-based structural analysis for the mutations. (**A**) and (**B**) Structures of spike-antibody binding interface shown as ribbons. Spike recognition of class III neutralizing antibodies C1365 (**A**) and SW186 (**B**) are interfered by R346S and K444T mutations, where multiple hydrogen bonds and salt-bridge (shown as yellow dot lines) are abolished. (**C**) Structure of spike-ACE2 binding interface shown as ribbon. F486 interacts hydrophobically with M82 and Y83 on ACE2, whereas F486S impedes this interaction. (**D**) Structural model of HR2 domain of SARS-CoV2 S. Inset: Electrostatic surface potential of HR2 membrane proximal region. D1199 contributes to the overall negative charge of this region.

## Discussion

We examined the neutralization resistance, fusogencitiy, and S processing of the emerging BA.4.6, BF.7, BQ.1, BQ.1.1, and BA.2.75.2 Omicron subvariants, as well as their lineage-defining mutations. All of these subvariants exhibit some degree of enhanced neutralization resistance over their parental BA.4/5 or BA.2.75.2 subvariants, with BQ.1, BQ.1.1, and BA.2.75.2 exhibiting the strongest resistance. Notably, this pattern is consistent for sera collected from HCWs following a homologous mRNA booster vaccination, from BA.1-wave hospitalized COVID-19 patients, as well as from BA.4/5-wave SARS-CoV-2 COVID-19 positive first responders and household contacts. Critically, the neutralization resistance of BQ.1 and BQ.1.1 appears to be driven largely by their N460K mutation, while the neutralization resistance of BA.2.75.2 variant is largely determined by the F486S mutation. Additionally, we provide evidence that the N460K and F486S mutations dictate the enhanced fusogenicity and S processing in their respective subvariants, with the D1199N mutation negatively modulating the fusogenicity of the BA.2.75.2 subvariant. Structural modeling and additional analyses revealed crucial roles of residues R436, K444 and F486 in antibody recognition and the potential mechanism of immune evasion through R436S, K444T and F486S mutations present in Omicron subvariants. Intriguingly, our structural analyses also suggest that D1199N mutation located in the HR2 region of S2 could alter the spike position in either an upright or a tilted status by reducing repulsion to the cellular membrane. A tilted spike with less efficient processing and receptor utilization could explain the reduced fusogenicity observed in D1199N-bearing mutations.

The neutralization resistance of the BQ.1, BQ.1.1, and BA.2.75.2 variants has concerning implications for the persistence of vaccine- and infection-induced immunity. The strong resistance of these variants to neutralization by patient sera, regardless of the immunogen — mRNA vaccination, BA.1 infection, or BA.4/5 infection — is particularly striking. This finding may indicate selection for immune evasion of even broadly neutralizing antibodies induced by multiple vaccinations and SARS-CoV-2 infections, as is now common in the population. In particular, the strong evasion of BA.4/5 infection-induced sera is concerning, as the recently recommended bivalent mRNA vaccine boosters contain BA.4/5 S along with the prototype. This possibility, together with the emergence of more diverse SARS-CoV-2 variants necessitates the development of more broadly active, even pan-coronavirus, COVID-19 vaccines (Chen et al., 2022; Su et al., 2022a). As circulating SARS-CoV-2 diversifies, the ability to vaccinate against dominant circulating variants may be even more compromised. Hence, examination of the efficacy of bivalent mRNA vaccines against emerging variants is critical to control and end the global pandemic.

In addition to changing the neutralization resistance, these new Omicron subvariants also affect the fusogenicity of the S proteins. Similar to our previous finding that BA.4/5 and BA.2.75 exhibit increased fusion capability (Qu et al., 2022a), here we also observed increased cell-cell fusion in several new Omicron subvariants compared to their respective parental subvariants, BA.4/5 or BA.2.75. These data may together indicate a continuing shift towards more efficient transmembrane protease, serine 2 (TMPRSS2) utilization to allow for plasma membrane-mediated viral fusion and entry pathway. Consistent with this notion, we observed an increased furin-cleavage efficiency in several S proteins, especially BQ.1, and BQ.1.1 subvariants, which could improve TMPRSS2 utilization (Mykytyn et al., 2021). This is concerning, because previously it has been shown that the shift in entry pathway from TMPRSS2-mediated plasma membrane fusion used by the prototype SARS-CoV-2 towards cathepsin B/L-mediated endosomal entry observed in the Omicron clade is associated with reduced lung tropism, increased nasopharyngeal tropism, and reduced pathogenicity of the Omicron variant (Barut et al., 2022; McMahan et al., 2022; Meng et al., 2022; Mykytyn et al., 2021; Shuai et al., 2022; Suzuki et al., 2022; Wang et al., 2022a). That said, here we show new Omicron subvariants continue to exhibit weak infection efficiency in lung-derived CaLu-3 cells. Examination of tissue tropism and pathogenicity *in vivo* for these and future emerging Omicron subvariants is critical to tailor appropriate public health responses.

In this work, we observed altered S processing and fusogenicity in Omicron subvariants. Notably, we found that many mutations examined, including N460K, N658S, and F486S, enhance S processing, as evidenced by increased S2/S ratios in viral producer cells. However, they fail to enhance, and in most cases reduce, the S1/S ratios in virus producer cells. This may indicate that the mutations likely destabilize the S1-S2 interaction and trimer conformation, consistent with increased cell-cell fusion and likely resulting enhanced S1 shedding. Intriguingly, we found that the D1199N mutation reduced S-mediated fusion and S processing, which could suggest that this mutation emerged to compensate for alterations in S functionality introduced by the F486S mutation. Further structural analyses on the impact of these key residues on S trimer conformation are needed to determine if these immune escape mutants are negatively impacting S functionality — potentially limiting the fitness of further immune escape mutations to be introduced.

The perpetual emergence of SARS-CoV-2 variants with enhanced immune escape continues to threaten public health. Monitoring the immune escape of emerging variants will be critical to improving mRNA vaccine reformulation, assessing new broader coronavirus vaccine candidates, as well as directing ongoing public health measures. Further, emerging variants must be monitored closely for any indication of selective pressure in enhancing lung tropism and potentially pathogenicity to ensure that any highly transmissible and more pathogenic variants are better and more quickly contained.

## Acknowledgements

We thank the NIH AIDS Reagent Program and BEI Resources for providing important reagents for this work. We also thank the Clinical Research Center/Center for Clinical Research Management of The Ohio State University Wexner Medical Center and The Ohio State University College of Medicine in Columbus, Ohio, specifically Francesca Madiai, Dina McGowan, Breona Edwards, Evan Long, and Trina Wemlinger, for logistics, collection and processing of samples. In addition, we thank Sarah Karow, Madison So, Preston So, Daniela Farkas, and Finny Johns in the clinical trials team of The Ohio State University for sample collection and other supports. This work was supported by a fund provided by an anonymous private donor to OSU. S.-L.L., F.S., D. J., G.L., A.P., R.J.G., L.J.S. and E.M.O. were supported by the National Cancer Institute of the NIH under award no. U54CA260582. The content is solely the responsibility of the authors and does not necessarily represent the official views of the National Institutes of Health. J.P.E. was supported by Glenn Barber Fellowship from the Ohio State University College of Veterinary Medicine. R.J.G. was additionally supported by the Robert J. Anthony Fund for Cardiovascular Research and the JB Cardiovascular Research Fund, and L.J.S. was partially supported by NIH R01 HD095881. K.X. was supported by the Ohio State University James Cancer Center and a Path to K award from the Ohio State University Office of Health Sciences and the Center for Clinical & Translational Science. The content is solely the responsibility of the authors and does not necessarily represent the official views of the university, or the Center for Clinical & Translational Science.

## Methods

### Samples and Patient Information

Sera samples were collected from HCWs at the Ohio State University Wexner Medical Center in Columbus, Ohio with approval from an institutional review board (Gordon et al.) (IRB 2020H0228 and IRB 2020H0527). These HCWs samples were collected 2-13 weeks after vaccination with a third homologous dose of the monovalent Moderna mRNA-1273 (n = 3) or Pfizer BioNTech BNT162b2 (n =12) vaccines. HCWs included 10 male and 5 female subjects with ages ranging from 26 to 61 (median 33).

Sera from BA.1-wave COVID-19 patients hospitalized in Columbus, Ohio were collected with approval from an IRB (IRB 2020H0527). The patient samples were collected 1-7 days after hospitalization with COVID-19. Hospitalizations occurred between the end of January and the end of February of 2022, a BA.1 dominant period in Columbus, Ohio. Patients included 12 male and 3 female patients with ages ranging from 29 to 78 (median 57). Patients included 6 unvaccinated patients, 5 patients vaccinated with 2 doses of Pfizer/BioNTech BNT162b2 (n = 2) or Moderna mRNA-1273 (n = 3), and 4 patients vaccinated and boosted with Pfizer/BioNTech BNT162b2.

Sera from BA.4/5-wave Columbus, Ohio first responders and household contacts who tested positive for SARS-CoV-2 infection were collected with IRB approval (IRB 2020H0527, 2020H0531, and 2020H0240). 11 patient nasal swab samples were sequenced to confirm infection with BA.4, BA.5, or a derivative variants, with 4 patients infected with BA.4, 7 with BA.5, and 9 patients could not have their variant determined. Of those who could not have the specific variant identified, their samples were collected between late July and late September of 2022, a BA.4/5 dominant period. Except for one patient whose gender and age are unknown, patients included 4 male and 15 female with ages ranging from 27 to 58 (median 44). Patients included 17 unvaccinated, and 3 vaccinated and boosted with Pfizer/BioNTech BNT162b2 (n=1) or Moderna mRNA-1273 (n = 2).

### Cell Lines and Maintenance

Human female embryonic kidney cell lines HEK293T (ATCC CRL-11268, RRID: CVCL_1926) and HEK293T cells stably expressing human ACE2 (BEI NR-52511, RRID: CVCL_A7UK) were maintained in DMEM (Gibco, 11965-092) with 10% FBS (Sigma, F1051) and 1% penicillin-streptomycin (HyClone, SV30010) added. Human male adenocarcinoma lung epithelial cell line Calu-3 (RRID:CVCL_0609) were maintained in EMEM (ATCC, 30-2003) with 10% FBS and 1% penicillin-streptomycin added. All cells were passaged first by washing with Dulbecco’s phosphate buffered saline (Sigma, D5652-10X1L) then incubating in 0.05% Trypsin + 0.53 mM EDTA (Corning, 25-052-CI) until cells were completely detached. Cells were maintained at 37°C and 5.0% CO2 in 10 cm cell culture dishes (Greiner Bio-one, 664160).

### Plasmids

The pNL4-3 inGluc lentiviral vector has been reported on in our previous publications (Zeng et al., 2020). Briefly, the vector is in the HIV-1 pNL4-3 backbone with a deletion of Env and an addition of a *Gaussia* luciferase reported gene that is expressed in target cells without premature expression in producer cells. The S variant constructs were cloned into the pcDNA3.1 vector by GenScript Biotech (Piscataway, NJ) using restriction enzyme cloning by KpnI and BamHI; alternatively, they were produced by PCR mutagenesis. The constructs bear FLAG tags on the N- and C-terminal ends. All constructs were confirmed by sequencing.

### Pseudotyped lentivirus production and infectivity

Pseudotyped lentiviral vectors were produced as previously described (Zeng et al., 2020). HEK293T cells were transfected with the pNL4-3-inGluc and S constructs in a 2:1 ratio using polyethyleneimine transfection (Transporter 5 Transfection Reagent, Polysciences) in order to generate viral particles. Virus was harvested 24, 48, and 72 hours post-transfection. Relative infectivity was determined by infection of HEK293T-ACE2 cells and measurement of *Gaussia* luciferase activity 48 hours post-infection. *Gaussia* luciferase activity was measured by combining equal volumes of cell culture media and *Gaussia* luciferase substrate (0.1 M Tris pH 7.4, 0.3 M sodium ascorbate, 10 μM coelenterazine) with luminescence measured immediately by a BioTek Cytation5 plate reader.

### Lentivirus neutralization assay

Neutralization assays with pseudotyped lentiviral vectors were performed as described previously (Zeng et al., 2020). HCW and COVID-19 patient samples were 4-fold serially diluted and equal amounts of SARS-CoV-2 pseudotyped virus were added to the diluted sera. Final dilutions were 1:80, 1:320, 1:1280, 1:5120, 1:20480, and no serum control. The virus and sera mixture was incubated for 1 hour at 37°C then added to HEK293T-ACE2 cells to allow for infection. *Gaussia* luciferase activity was measured at 48 and 72 hours post-infection by combining equal volumes of cell culture media and *Gaussia* luciferase substrate with luminescence measured immediately by a BioTek Cytation5 plate reader. The 50% neutralization titers (NT50) were determined by least-squares-fit, non-linear regression in GraphPad Prism 9 (San Diego, CA).

### Spike surface expression

HEK293T cells used to produce pseudotyped lentiviral vectors were harvested 72 hours post-transfection. The producer cells were incubated in PBS+5mM EDTA for 30 minutes at 37C to disassociate them. The cells were then fixed in 4% formaldehyde and stained with anti-SARS-CoV-2 polyclonal S1 antibody (Sino Biological, 40591-T62; RRID: AB_2893171). Cells were then stained with anti-rabbit-IgG-FITC (Sigma, F9887, RRID: AB_259816) and assayed with a Life Technologies Attune NxT flow cytometer. FlowJo v7.6.5 (Ashalnd, OR) was used to process flow cytometry data.

### Syncytia formation

HEK293T-ACE2 cells were co-transfected with the S of interest alongside GFP. Then 24 hours post transfection, the cells were imaged on Leica DMi8 confocal microscope to visualize syncytia. Representative images were selected. Mean syncytia size was determine using Leica X Applications Suite. The scale bars represent 150 μm.

### Spike processing and incorporation

Lysate was collected from virus producer cells through a 30-minute incubation on ice in RIPA lysis buffer (50 mM Tris pH 7.5, 150 mM NaCl, 1 mM EDTA, Nonidet P-40, 0.1% SDS) supplemented with protease inhibitor (Sigma, P8340). Samples were run on a 10% acrylamide SDS-PAGE gel and transferred to a PVDF membrane. Membranes were probed with anti-S1 (Sino Biological, 40591-T62; RRID:AB_2893171), anti-S2 (Sino Biological, 40590; RRID:AB_2857932), and anti-β-actin (ThermoFisher, MA5-15740; RRID:AB_10983927). Secondary antibodies included Anti-mouse-IgG-Peroxidase (Sigma, A5278; RRID:AB_258232) and Anti-rabbit-IgG-HRP (Sigma, A9169; RRID:AB_258434). Blots were imaged using Immobilon Crescendo Western HRP substrate (Millipore, WBLUR0500) on a GE Amersham Imager 600. Band intensities were quantified using NIH ImageJ (Bethesda, MD) image analysis software.

### Structural modeling and analysis

Homology modeling of Omicron spike protein complexes with either ACE2 receptor or neutralizing antibodies was performed on SWISS-MODEL server with published X-ray crystallography and cryo-EM structures as templates (PDB IDs: 7K8Z, 8DT3, 7XB0, 2FXP). Molecular contacts of Omicron mutants were examined and illustrated with the programs PyMOL.

### Quantification and statistical analysis

NT50 values were determined by least-squares-fit, non-linear-regression in GraphPad Prism 9 (San Diego, CA). NT_50_ values were log_10_ transformed for hypothesis testing to better approximate normality, and multiplicity was addressed by the use of Bonferroni corrections. The statistical analysis was performed using GraphPad Prism 9 and are referenced in the figure legends, including one-way ANOVA (Figs. 1B-C, 2A-B, 3B, 3D, 4B, 4D, 4E, 4F, and S2A-B) and one-way repeated measures ANOVA (Figs. 1D–F, and 2C-H). Syncytia sizes (Figs. 3B, 4B, and 4D) were quantified by Leica Applications Suit X (Wetzlar, Germany). Band intensities (Figures 3E, 4G, and 4H) were quantified using NIH ImageJ (Bethesda, MD) analysis software.

## References

Abu-Raddad, L.J., Chemaitelly, H., Ayoub, H.H., AlMukdad, S., Yassine, H.M., Al-Khatib, H.A., Smatti, M.K., Tang, P., Hasan, M.R., Coyle, P., et al. (2022). Effect of mRNA Vaccine Boosters against SARS-CoV-2 Omicron Infection in Qatar. The New England journal of medicine 386, 1804–1816.

Barut, G.T., Halwe, N.J., Taddeo, A., Kelly, J.N., Schön, J., Ebert, N., Ulrich, L., Devisme, C., Steiner, S., Trüeb, B.S., et al. (2022). The spike gene is a major determinant for the SARS-CoV-2 Omicron-BA.1 phenotype. Nat Commun 13, 5929.

Cao, Y., Song, W., Wang, L., Liu, P., Yue, C., Jian, F., Yu, Y., Yisimayi, A., Wang, P., Wang, Y., et al. (2022a). Characterization of the enhanced infectivity and antibody evasion of Omicron BA. 2.75. Cell Host & Microbe.

Cao, Y., Wang, J., Jian, F., Xiao, T., Song, W., Yisimayi, A., Huang, W., Li, Q., Wang, P., An, R., et al. (2022b). Omicron escapes the majority of existing SARS-CoV-2 neutralizing antibodies. Nature 602, 657–663.

Cao, Y., Yisimayi, A., Jian, F., Song, W., Xiao, T., Wang, L., Du, S., Wang, J., Li, Q., Chen, X., et al. (2022c). BA.2.12.1, BA.4 and BA.5 escape antibodies elicited by Omicron infection. Nature 608, 593–602.

Centers for Disease Control and Prevention (2022). COVID Data Tracker. Atlanta, GA: US Department of Health and Human Services, CDC; 2022, October 16. https://covid.cdc.gov/covid-data-tracker.

Cerutti, G., Guo, Y., Liu, L., Liu, L., Zhang, Z., Luo, Y., Huang, Y., Wang, H.H., Ho, D.D., Sheng, Z., et al. (2022). Cryo-EM structure of the SARS-CoV-2 Omicron spike. Cell Rep 38, 110428.

Chen, Y., Zhao, X., Zhou, H., Zhu, H., Jiang, S., and Wang, P. (2022). Broadly neutralizing antibodies to SARS-CoV-2 and other human coronaviruses. Nature reviews Immunology, 1–11.

Cui, Z., Liu, P., Wang, N., Wang, L., Fan, K., Zhu, Q., Wang, K., Chen, R., Feng, R., Jia, Z., et al. (2022). Structural and functional characterizations of infectivity and immune evasion of SARS-CoV-2 Omicron. Cell 185, 860–871.e813.

Evans, J.P., Zeng, C., Qu, P., Faraone, J., Zheng, Y.M., Carlin, C., Bednash, J.S., Zhou, T., Lozanski, G., Mallampalli, R., et al. (2022). Neutralization of SARS-CoV-2 Omicron sub-lineages BA.1, BA.1.1, and BA.2. Cell Host Microbe 30, 1093–1102.e1093.

Gordon, D.E., Jang, G.M., Bouhaddou, M., Xu, J., Obernier, K., White, K.M., O’Meara, M.J., Rezelj, V.V., Guo, J.Z., Swaney, D.L., et al. (2020). A SARS-CoV-2 protein interaction map reveals targets for drug repurposing. Nature 583, 459–468.

Gruell, H., Vanshylla, K., Tober-Lau, P., Hillus, D., Schommers, P., Lehmann, C., Kurth, F., Sander, L.E., and Klein, F. (2022). mRNA booster immunization elicits potent neutralizing serum activity against the SARS-CoV-2 Omicron variant. Nat Med 28, 477–480.

Hachmann, N.P., Miller, J., Collier, A.Y., Ventura, J.D., Yu, J., Rowe, M., Bondzie, E.A., Powers, O., Surve, N., Hall, K., et al. (2022). Neutralization Escape by SARS-CoV-2 Omicron Subvariants BA.2.12.1, BA.4, and BA.5. The New England journal of medicine 387, 86–88.

Iacobucci, G. (2022). Covid-19: Hospital admissions rise in England as some trusts reinstate mask requirements. Bmj 379, o2440.

Iketani, S., Liu, L., Guo, Y., Liu, L., Chan, J.F., Huang, Y., Wang, M., Luo, Y., Yu, J., Chu, H., et al. (2022). Antibody evasion properties of SARS-CoV-2 Omicron sublineages. Nature 604, 553–556.

Khan, K., Karim, F., Ganga, Y., Bernstein, M., Jule, Z., Reedoy, K., Cele, S., Lustig, G., Amoako, D., Wolter, N., et al. (2022). Omicron BA.4/BA.5 escape neutralizing immunity elicited by BA.1 infection. Nat Commun 13, 4686.

Kimura, I., Yamasoba, D., Tamura, T., Nao, N., Suzuki, T., Oda, Y., Mitoma, S., Ito, J., Nasser, H., Zahradnik, J., et al. (2022). Virological characteristics of the SARS-CoV-2 Omicron BA.2 subvariants, including BA.4 and BA.5. Cell 185, 3992–4007.e3916.

Kurhade, C., Zou, J., Xia, H., Cai, H., Yang, Q., Cutler, M., Cooper, D., Muik, A., Jansen, K.U., Xie, X., et al. (2022). Neutralization of Omicron BA.1, BA.2, and BA.3 SARS-CoV-2 by 3 doses of BNT162b2 vaccine. Nat Commun 13, 3602.

Liu, L., Iketani, S., Guo, Y., Chan, J.F., Wang, M., Liu, L., Luo, Y., Chu, H., Huang, Y., Nair, M.S., et al. (2022). Striking antibody evasion manifested by the Omicron variant of SARS-CoV-2. Nature 602, 676–681.

Lyngse, F.P., Kirkeby, C.T., Denwood, M., Christiansen, L.E., Mølbak, K., Møller, C.H., Skov, R.L., Krause, T.G., Rasmussen, M., Sieber, R.N., et al. (2022). Transmission of sars-cov-2 omicron voc subvariants BA.1 and BA.2: Evidence from danish households. MedRxiv.

McMahan, K., Giffin, V., Tostanoski, L.H., Chung, B., Siamatu, M., Suthar, M.S., Halfmann, P., Kawaoka, Y., Piedra-Mora, C., Jain, N., et al. (2022). Reduced pathogenicity of the SARS-CoV-2 omicron variant in hamsters. Med (New York, NY) 3, 262–268.e264.

Meng, B., Abdullahi, A., Ferreira, I., Goonawardane, N., Saito, A., Kimura, I., Yamasoba, D., Gerber, P.P., Fatihi, S., Rathore, S., et al. (2022). Altered TMPRSS2 usage by SARS-CoV-2 Omicron impacts infectivity and fusogenicity. Nature 603, 706–714.

Mykytyn, A.Z., Breugem, T.I., Riesebosch, S., Schipper, D., van den Doel, P.B., Rottier, R.J., Lamers, M.M., and Haagmans, B.L. (2021). SARS-CoV-2 entry into human airway organoids is serine protease-mediated and facilitated by the multibasic cleavage site. eLife 10.

Peacock, T.P., Goldhill, D.H., Zhou, J., Baillon, L., Frise, R., Swann, O.C., Kugathasan, R., Penn, R., Brown, J.C., Sanchez-David, R.Y., et al. (2021). The furin cleavage site in the SARS-CoV-2 spike protein is required for transmission in ferrets. Nat Microbiol 6, 899–909.

Qu, P., Evans, J.P., Zheng, Y.M., Carlin, C., Saif, L.J., Oltz, E.M., Xu, K., Gumina, R.J., and Liu, S.L. (2022a). Evasion of neutralizing antibody responses by the SARS-CoV-2 BA.2.75 variant. Cell Host Microbe.

Qu, P., Faraone, J., Evans, J.P., Zou, X., Zheng, Y.M., Carlin, C., Bednash, J.S., Lozanski, G., Mallampalli, R.K., Saif, L.J., et al. (2022b). Neutralization of the SARS-CoV-2 Omicron BA.4/5 and BA.2.12.1 Subvariants. N Engl J Med 386, 2526–2528.

Saito, A., Tamura, T., Zahradnik, J., Deguchi, S., Tabata, K., Anraku, Y., Kimura, I., Ito, J., Yamasoba, D., Nasser, H., et al. Virological characteristics of the SARS-CoV-2 Omicron BA. 2.75 variant. Cell Host & Microbe.

Shuai, H., Chan, J.F., Hu, B., Chai, Y., Yuen, T.T., Yin, F., Huang, X., Yoon, C., Hu, J.C., Liu, H., et al. (2022). Attenuated replication and pathogenicity of SARS-CoV-2 B.1.1.529 Omicron. Nature 603, 693–699.

Stegger, M., Edslev, S.M., Sieber, R.N., Ingham, A.C., Ng, K.L., Tang, M.-H.E., Alexandersen, S., Fonager, J., Legarth, R., Utko, M., et al. (2022). Occurrence and significance of Omicron BA. 1 infection followed by BA. 2 reinfection. medRxiv.

Su, S., Li, W., and Jiang, S. (2022a). Developing pan-β-coronavirus vaccines against emerging SARS-CoV-2 variants of concern. Trends Immunol 43, 170–172.

Su, W., Choy, K.T., Gu, H., Sia, S.F., Cheng, K.M., Nizami, S.I.N., Krishnan, P., Ng, Y.M., Chang, L.D.J., Liu, Y., et al. (2022b). Omicron BA.1 and BA.2 sub-lineages show reduced pathogenicity and transmission potential than the early SARS-CoV-2 D614G variant in Syrian hamsters. J Infect Dis.

Suzuki, R., Yamasoba, D., Kimura, I., Wang, L., Kishimoto, M., Ito, J., Morioka, Y., Nao, N., Nasser, H., Uriu, K., et al. (2022). Attenuated fusogenicity and pathogenicity of SARS-CoV-2 Omicron variant. Nature 603, 700–705.

Tuekprakhon, A., Nutalai, R., Dijokaite-Guraliuc, A., Zhou, D., Ginn, H.M., Selvaraj, M., Liu, C., Mentzer, A.J., Supasa, P., Duyvesteyn, H.M.E., et al. (2022). Antibody escape of SARS-CoV-2 Omicron BA.4 and BA.5 from vaccine and BA.1 serum. Cell 185, 2422–2433.e2413.

Wang, Q., Anang, S., Iketani, S., Guo, Y., Liu, L., Katsamba, P.S., Shapiro, L., Ho, D.D., and Sodroski, J.G. (2022a). Functional properties of the spike glycoprotein of the emerging SARS-CoV-2 variant B.1.1.529. Cell Rep 39, 110924.

Wang, Q., Guo, Y., Iketani, S., Nair, M.S., Li, Z., Mohri, H., Wang, M., Yu, J., Bowen, A.D., Chang, J.Y., et al. (2022b). Antibody evasion by SARS-CoV-2 Omicron subvariants BA.2.12.1, BA.4 and BA.5. Nature 608, 603–608.

Wang, Q., Iketani, S., Li, Z., Guo, Y., Yeh, A.Y., Liu, M., Yu, J., Sheng, Z., Huang, Y., Liu, L., et al. (2022c). Antigenic characterization of the SARS-CoV-2 Omicron subvariant BA.2.75. Cell Host Microbe.

Xia, H., Zou, J., Kurhade, C., Cai, H., Yang, Q., Cutler, M., Cooper, D., Muik, A., Jansen, K.U., Xie, X., et al. (2022a). Neutralization and durability of 2 or 3 doses of the BNT162b2 vaccine against Omicron SARS-CoV-2. Cell Host Microbe 30, 485–488.e483.

Xia, S., Wang, L., Zhu, Y., Lu, L., and Jiang, S. (2022b). Origin, virological features, immune evasion and intervention of SARS-CoV-2 Omicron sublineages. Signal Transduct Target Ther 7, 241.

Yamasoba, D., Kimura, I., Nasser, H., Morioka, Y., Nao, N., Ito, J., Uriu, K., Tsuda, M., Zahradnik, J., Shirakawa, K., et al. (2022). Virological characteristics of the SARS-CoV-2 Omicron BA.2 spike. Cell 185, 2103–2115.e2119.

Yu, J., Collier, A.Y., Rowe, M., Mardas, F., Ventura, J.D., Wan, H., Miller, J., Powers, O., Chung, B., Siamatu, M., et al. (2022). Neutralization of the SARS-CoV-2 Omicron BA.1 and BA.2 Variants. The New England journal of medicine 386, 1579–1580.

Zeng, C., Evans, J.P., Pearson, R., Qu, P., Zheng, Y.M., Robinson, R.T., Hall-Stoodley, L., Yount, J., Pannu, S., Mallampalli, R.K., et al. (2020). Neutralizing antibody against SARS-CoV-2 spike in COVID-19 patients, health care workers, and convalescent plasma donors. JCI Insight 5.

Zeng, C., Evans, J.P., Qu, P., Faraone, J., Zheng, Y.M., Carlin, C., Bednash, J.S., Zhou, T., Lozanski, G., Mallampalli, R., et al. (2021). Neutralization and Stability of SARS-CoV-2 Omicron Variant. bioRxiv.

Zou, J., Xia, H., Xie, X., Kurhade, C., Machado, R.R.G., Weaver, S.C., Ren, P., and Shi, P. Y. (2022). Neutralization against Omicron SARS-CoV-2 from previous non-Omicron infection. Nat Commun 13, 852.

