## Supplemental Appendix for "Distinct Neutralizing Antibody Escape of SARS-CoV-2 Omicron Subvariants BQ.1, BQ.1.1, BA.4.6, BF.7 and BA.2.75.2"

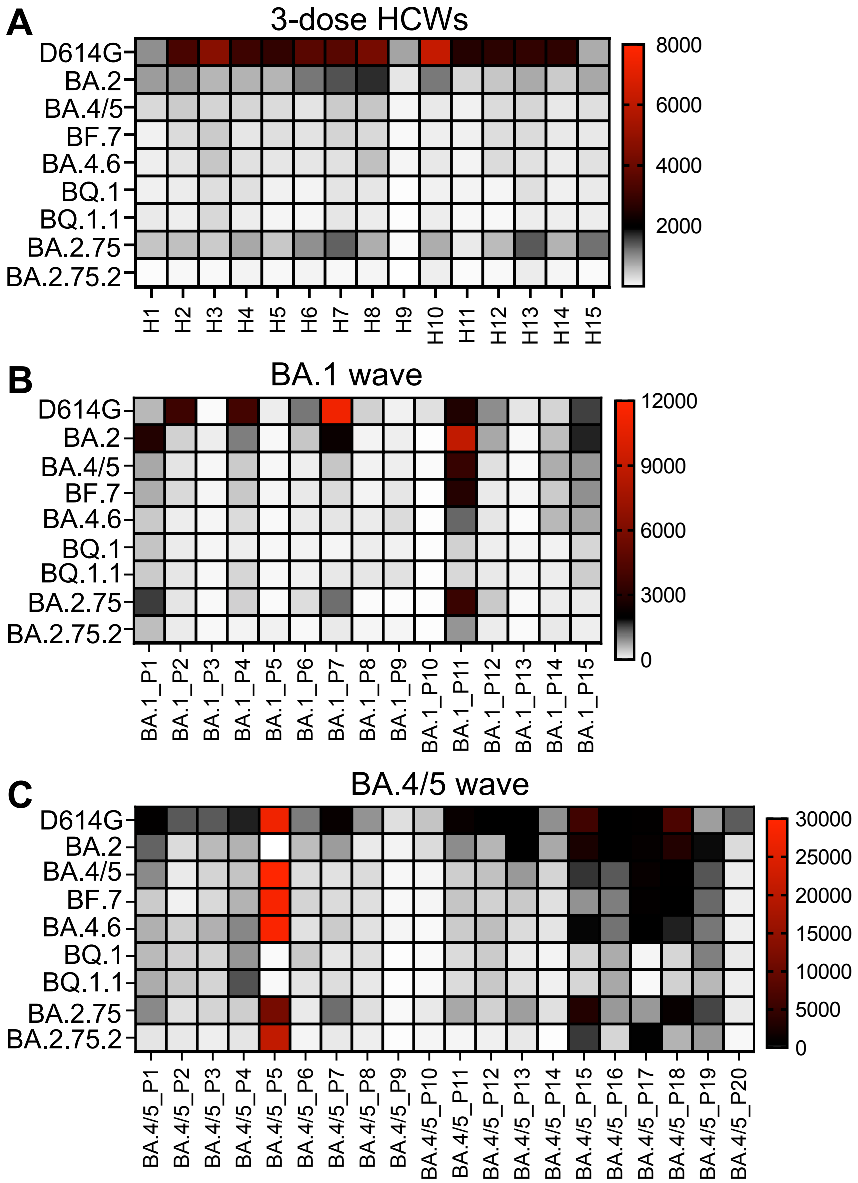


**Figure S1: Omicron-derived subvariants display distinct immune escape pattens.**
(**A-C**) Neutralizing antibody titers are displayed as a heat map for sera from health care workers (HCWs, “H#”) (n=15) who received a single homologous monovalent Moderna mRNA-1273 (n =3) or Pfizer/BioNTech BNT162b2 (n =12) mRNA booster vaccination (A), for sera from BA.1-wave hospitalized COVID-19 patients (“BA.1_P#”) (n = 15) (B), and for sera from BA.4/5-wave SARS-CoV-2 infected Columbus, Ohio first responders and household contacts (“BA.4/5_P#”) (n = 20) (C).

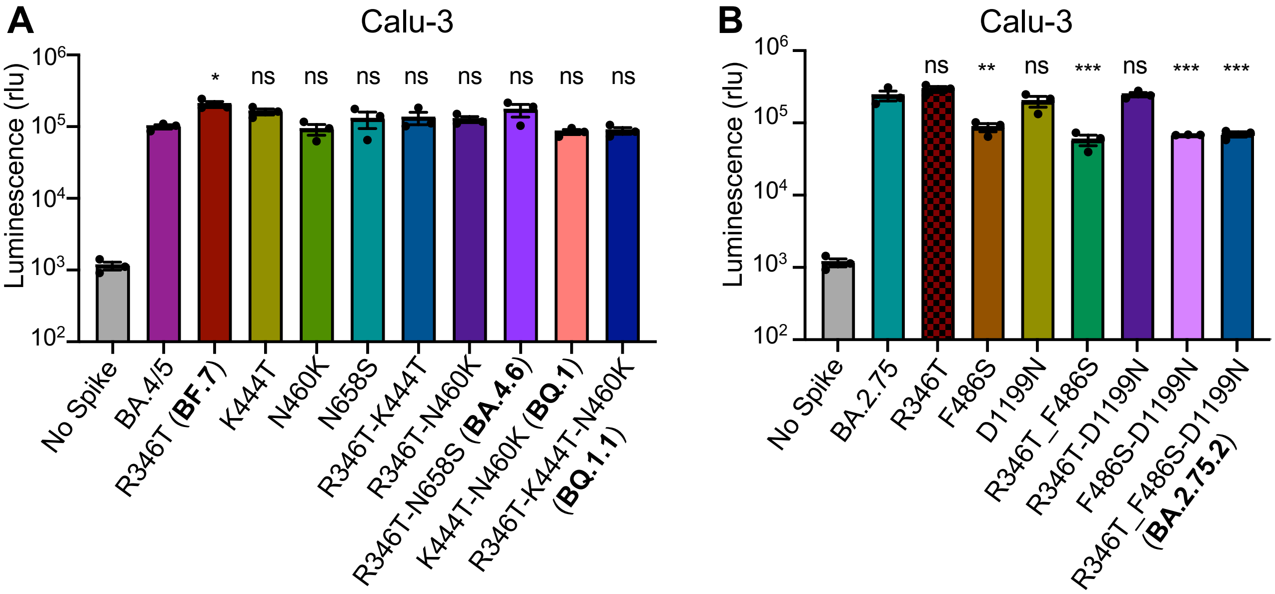


**Figure S2: Infectivity of BA.4/5- and BA.2.75-derived mutants in Calu-3 cells.**
(**A-B**) Infectivity of lentivirus pseudotyped with the indicated BA.4/5-derived mutant S constructs, n = 3 (A) or BA.2.75-derived mutant S constructs, n = 3 (B) in CaLu-3 cells. Bars represent means +/- standard error. Significance relative to BA.4/5 or BA.2.75 was determined by one-way ANOVA with Bonferroni's multiple testing correction. P-values are represented as ns for $p \geq 0.05$, *$p < 0.05$, **$p < 0.01$, and ***$p < 0.001$.

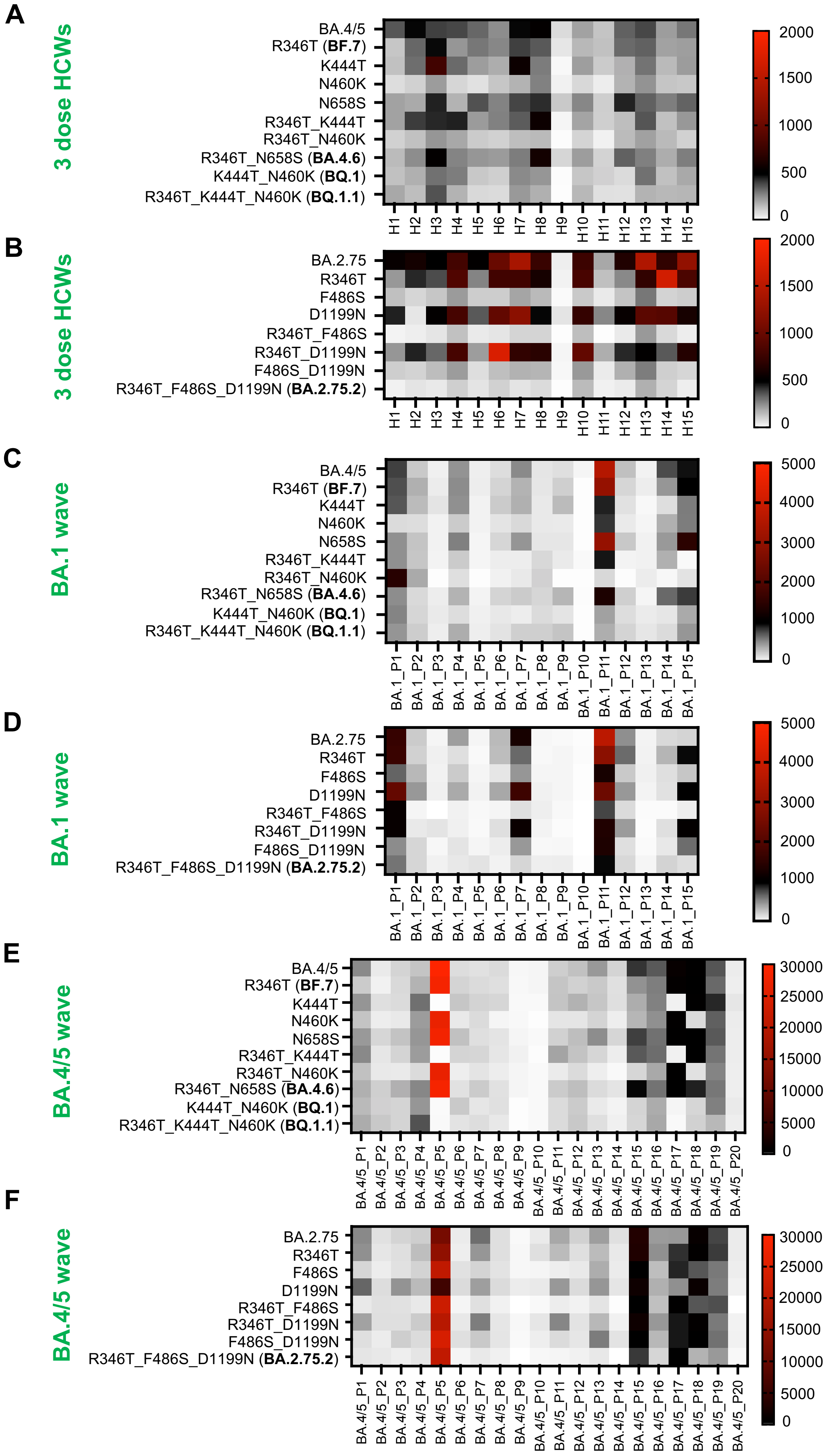

A
3 dose HCWs
BA.4/5
R346T (BF.7)
K444T
N460K
N658S
R346T_K444T
R346T_N460K
R346T_N658S (BA.4.6)
K444T_N460K (BQ.1)
R346T_K444T_N460K (BQ.1.1)
H1 H2 H3 H4 H5 H6 H7 H8 H9 H10 H11 H12 H13 H14 H15
2000 1500 1000 500 0
B
3 dose HCWs
BA.2.75
R346T
F486S
D1199N
R346T_F486S
R346T_D1199N
F486S_D1199N
R346T_F486S_D1199N (BA.2.75.2)
H1 H2 H3 H4 H5 H6 H7 H8 H9 H10 H11 H12 H13 H14 H15
2000 1500 1000 500 0
C
BA.1 wave
BA.4/5
R346T (BF.7)
K444T
N460K
N658S
R346T_K444T
R346T_N460K
R346T_N658S (BA.4.6)
K444T_N460K (BQ.1)
R346T_K444T_N460K (BQ.1.1)
BA.1_P1 BA.1_P2 BA.1_P3 BA.1_P4 BA.1_P5 BA.1_P6 BA.1_P7 BA.1_P8 BA.1_P9 BA.1_P10 BA.1_P11 BA.1_P12 BA.1_P13 BA.1_P14 BA.1_P15
5000 4000 3000 2000 1000 0
D
BA.1 wave
BA.2.75
R346T
F486S
D1199N
R346T_F486S
R346T_D1199N
F486S_D1199N
R346T_F486S_D1199N (BA.2.75.2)
BA.1_P1 BA.1_P2 BA.1_P3 BA.1_P4 BA.1_P5 BA.1_P6 BA.1_P7 BA.1_P8 BA.1_P9 BA.1_P10 BA.1_P11 BA.1_P12 BA.1_P13 BA.1_P14 BA.1_P15
5000 4000 3000 2000 1000 0
E
BA.4/5 wave
BA.4/5
R346T (BF.7)
K444T
N460K
N658S
R346T_K444T
R346T_N460K
R346T_N658S (BA.4.6)
K444T_N460K (BQ.1)
R346T_K444T_N460K (BQ.1.1)
BA.4/5_P1 BA.4/5_P2 BA.4/5_P3 BA.4/5_P4 BA.4/5_P5 BA.4/5_P6 BA.4/5_P7 BA.4/5_P8 BA.4/5_P9 BA.4/5_P10 BA.4/5_P11 BA.4/5_P12 BA.4/5_P13 BA.4/5_P14 BA.4/5_P15 BA.4/5_P16 BA.4/5_P17 BA.4/5_P18 BA.4/5_P19 BA.4/5_P20
30000 25000 20000 15000 10000 5000 0
F
BA.4/5 wave
BA.2.75
R346T
F486S
D1199N
R346T_F486S
R346T_D1199N
F486S_D1199N
R346T_F486S_D1199N (BA.2.75.2)
BA.4/5_P1 BA.4/5_P2 BA.4/5_P3 BA.4/5_P4 BA.4/5_P5 BA.4/5_P6 BA.4/5_P7 BA.4/5_P8 BA.4/5_P9 BA.4/5_P10 BA.4/5_P11 BA.4/5_P12 BA.4/5_P13 BA.4/5_P14 BA.4/5_P15 BA.4/5_P16 BA.4/5_P17 BA.4/5_P18 BA.4/5_P19 BA.4/5_P20
30000 25000 20000 15000 10000 5000 0


Figure S3

**Figure S3: BA.4/5- and BA.2.75-derived mutants display distrinct immune escapes.**
(**A-F**) Neutralizing antibody tiers against BA.4/5-derived (A, C and E) or BA.2.75-derived (B, D and F) mutants are displayed as heat maps for sera from health care workers (HCWs, "H#") (n = 15) who received a single homologous monovalent Moderna mRNA-1273 (n = 3) or Pfizer/BioNTech BNT162b2 (n = 12) mRNA booster vaccination (A, B); for sera from BA.1-wave hospitalized COVID-19 patients ("BA.1_P#") (n = 15) (C, D), and for sera from BA.4/5-wave SARS-CoV-2 infected Columbus, Ohio first responders and household contacts ("BA.4/5_P#") (n = 20) (E and F).

Figure S3

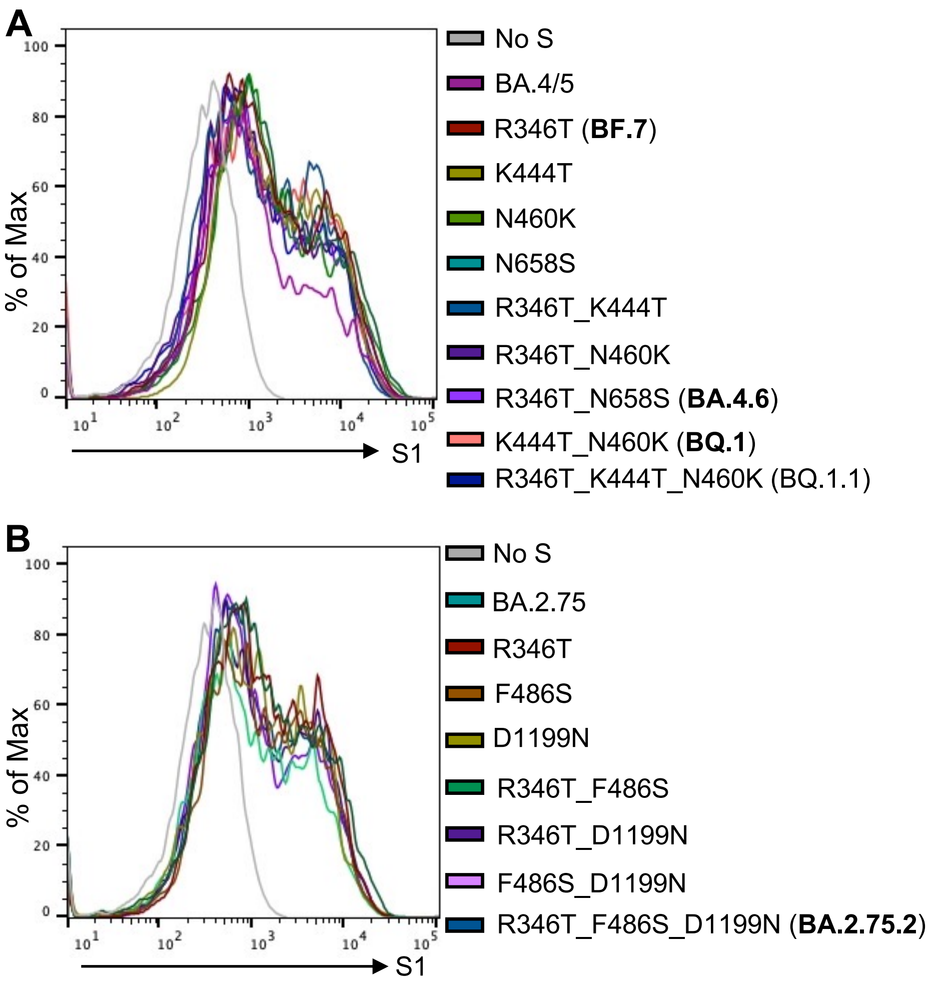


**Figure S4: Surface expression of BA.4/5 and BA.2.75 derived mutants.**
(**A-B**) Representative histograms of anti-S surface staining in HEK293T cells transfect to express BA.4/5 derived mutant S proteins (A) or BA.2.75 derived mutant S proteins (B).
